# Automated Eight-Stage Classification of *Drosophila melanogaster* Using Transfer-Learning CNNs with Mobile Live-Inference Deployment

**DOI:** 10.1101/2025.09.19.677285

**Authors:** Ridhee Bhatt, Rishi Prasad, S Aswini, Soumyadeep Paul, Ishaan Gupta

**Author notes:** Correspondence to Ishaan Gupta. Co-first author (Ridhee Bhatt, Rishi Prasad).

## Abstract

*Drosophila melanogaster* is a foundational model organism whose rapid development and genetic tractability underpin research in genetics, development, and disease. Manual staging of its embryonic, larval, and pupal phases is slow and error-prone. An automated eight-class classifier is introduced to distinguish egg, first-, second-, and third-instar larvae, as well as white, brown, eye, and black pupae, from stereo-microscope images. By fine-tuning ImageNet-pretrained CNNs (ResNet-50, InceptionV3, ConvNeXtSmall) on a balanced dataset (∼300 images per class), the best-performing model (ResNet-50) achieves 85% accuracy (F1 = 0.85) on a held-out validation set, significantly outperformed alternatives. Primary misclassifications align with subtle morphological transitions between adjacent stages. To facilitate broad adoption, the ResNet-50 model has been deployed in a lightweight Streamlit app offering live-camera inference (≈8 FPS on mobile). All code, pretrained weights, and data are publicly available, enabling scalable, high-throughput *Drosophila* staging for diverse experimental workflows.

## INTRODUCTION

*Drosophila melanogaster*, also known as the fruit fly, is a well-studied model organism that has played a pivotal role in advancing biological domains, including genetics, developmental biology, and disease modeling. This fruit fly, although morphologically very distinct from humans, shares over 60% of human disease-associated genes with us ^1,2^. Moreover, its short generation time, simple genetic structure, and ease of laboratory maintenance make it a model organism, one that has been used for over a century. Although the reason as to why *Drosophila* was first used isn’t known, its rise to prominence is well-documented ^3^. The sex limited inheritance experiment performed by Thomas Hunt Morgan proved to be a turning point not only for modern genetics but also for *Drosophila* as a model organism ^4^.

The life cycle of *Drosophila* is divided into four major stages: egg, larva, pupa, and adult^1^. The entire life cycle spans around 10 days at 25°C. *Drosophila*, being a holometabolous insect, undergoes complete metamorphosis during its life cycle. The first stage, i.e., the egg, is white, oval-shaped, with tapered ends. It has two respiratory filaments at the anterior end. The first instar has a translucent white colour and lacks strong segmentation. Its mouth hooks are barely visible. The second instar, however, is more robust than the first instar. It is slightly opaque white in colour and shows clearer segmentation on its body. The third instar is relatively the most active stage out of the three instars. It is found on the sides of the vial and is ready to pupate. It is thick, cylindrical with tapered ends, and whitish-cream to pale yellow in colour. It has distinct segmentation, visible black mouth hooks, and higher fat body content.

The pupal stages have been divided into four major stages on the basis of visual developmental features: white pupa, brown pupa, eye pupa, and black pupa. The white pupae is a short transitional phase, also called as the pre-pupal stage, where the third instar immobilizes and starts to pupate. It is an early stage of pupa. As the pupa develops, it obtains a brownish body with minimal to no body segmentation. This is the brown pupa stage. This stage then develops into the eye pupa stage, where reddish-brown spots appear at the location of adult eyes. The last stage, i.e. the black pupa, has a darker body with well-defined segmentation and visible adult structures like compound eyes, bristles, legs, wings, and genitalia.

By examining the research applications of each stage, one can understand how a single genome can orchestrate the development of multiple phenotypes, adhering to the principles of biology. The embryonic stage is a cornerstone of developmental biology, as it provides a remarkably clear and genetically tractable system for investigating how a complex body plan is established from a simple set of initial instructions. The principles of axis formation, segmentation, and cell fate determination, first elucidated in *Drosophila* embryos, have proven to be fundamental to the development of all bilaterally symmetric animals, including humans ^5^.

The larval stage is a critical developmental window during which the precursors of the adult body, the imaginal discs, proliferate and are patterned. The study of the larval stage reveals that it is far more than a simple growth phase. It is a period of developmental programming modulated by external cues, particularly nutrition ^6^. A hallmark of metamorphosis is the systematic destruction of most larval-specific tissues, in the pupal stage, through a process called histolysis^7^. With the destruction of the larval tissues, the adult body starts forming. The study of pupal development offers profound insights into the logic of hormonal control ^7^.

Manual classification of *Drosophila* life stages is a time-consuming and labor-intensive process that requires expert precision. In the age of artificial intelligence, this task can be automated using supervised learning techniques ^8^. This work presents deep learning models trained to classify *Drosophila melanogaster* across eight developmental stages: egg, first instar, second instar, third instar, white pupa, brown pupa, eye pupa, and black pupa.

Each class included approximately 300 annotated images. Convolutional neural networks (CNNs) ^9^ were used, including ResNet50 ^10^, InceptionV3 ^11^ and ConvNeXt ^12^, leveraging transfer learning to fine-tune these models for the specific classification tasks. The trained models were then integrated into a lightweight, user-friendly web application that supports both image uploads and live camera-based predictions. Looking at the results section, it can be seen that the ResNet50 model provides the highest accuracy of 85%.

## METHODOLOGY

### Imaging Protocol for Developmental Stage Classification in *Drosophila melanogaster*

The Oregon-R strain of *Drosophila* was used for image collection. This strain is widely used in genetic, developmental biology, and neurobiology research due to its robust health and high fertility ^8^.

To collect images of different life stages of *Drosophila melanogaster*, a tightly timed and controlled rearing protocol was followed to obtain the population. Firstly, adult flies were placed in fresh sterile vials with cornstarch media. They were then kept at 25 ± 1°C under a 12-hour light/dark cycle. Eggs were collected the next day, such that all flies were at the same developmental stage. This time was marked as 0 hours after egg laying (AEL). Images of the eggs were taken immediately.

Larvae were collected and imaged at regular time points: 24 hours AEL (1st instar), 48 hours AEL (2nd instar), and 72–96 hours AEL (3rd instar). Larvae were gently washed in PBS and placed on the stage plate for imaging. Pupae were also photographed starting at 96 hours AEL, including both early and late pupal stages. All images were taken using a T60N Series Stereo Microscope with a digital camera under consistent lighting and background (Figure:1).

**Figure 1.**
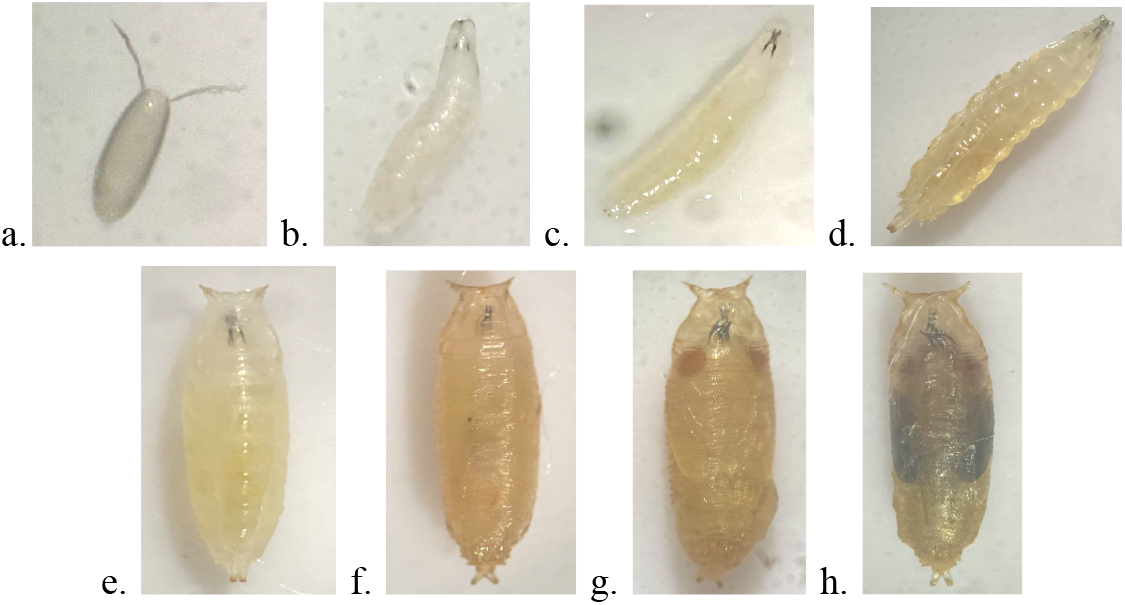
Developmental stages of *Drosophila melanogaster* showing progressive changes in morphology: a. Egg, b. First Instar, c. Second Instar, d. Third Instar, e. White Pupa, f. Brown Pupa, g. Eye Pupa, and h. Black Pupa

**Figure 2.**
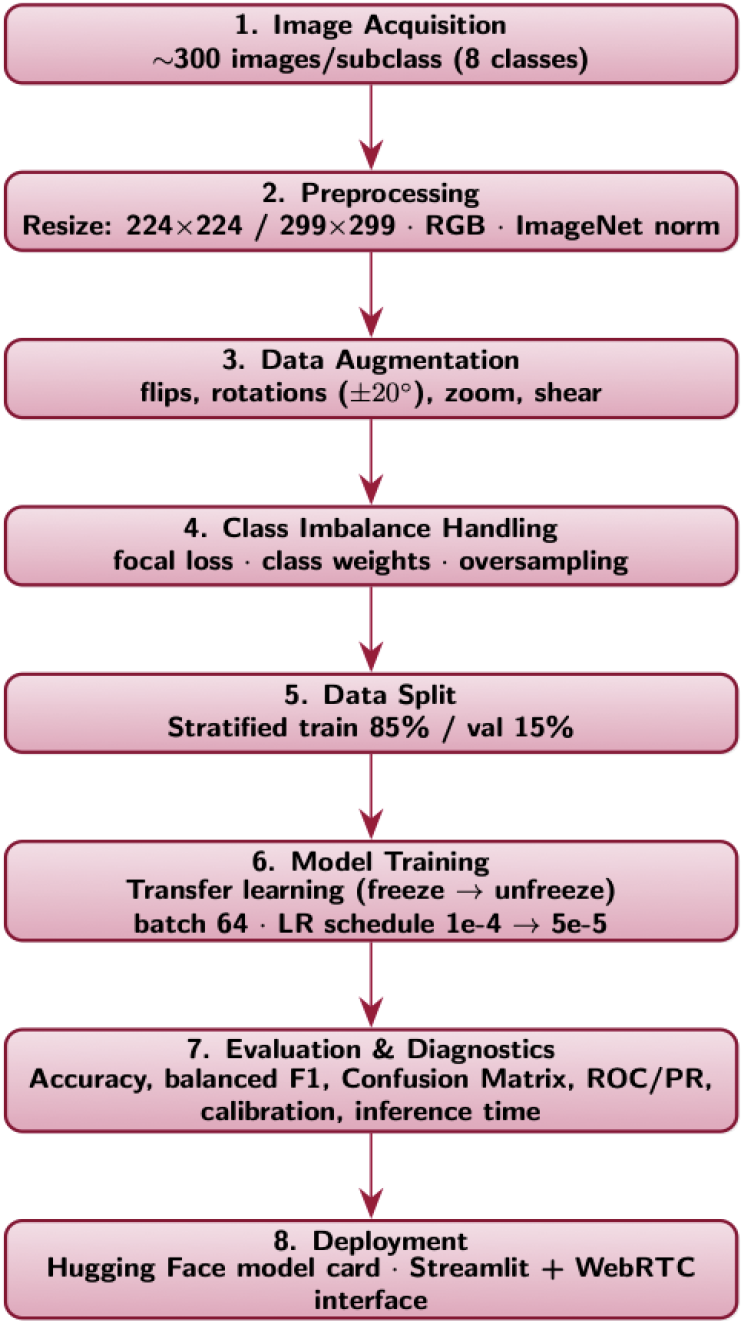
End-to-end workflow for *Drosophila* life stage classification

### Deep Learning Pipeline for Robust *Drosophila* Stage Classification

#### From Rule Based Processing to CNN

For the classification process, rather than relying on rule-based models, convolutional neural networks (CNNs) were used for the development of models. Traditional rule-based image processing uses hand-crafted algorithms like thresholding and filters, which perform well in controlled conditions but often fail under variations in lighting, pose, or background. In contrast, CNNs automatically learn features at multiple levels, making them more robust and adaptable to real-world variability with minimal manual tuning.

#### Image Preprocessing and data augmentation

Before feeding the *Drosophila* photographs into any neural network, a systematic preprocessing pipeline was performed to ensure consistency, augment diversity, and accelerate convergence. First, every raw image—whether captured under slightly different lighting or magnification—was resized to a fixed dimension (224×224 for ResNet50, 299×299 for InceptionV3) and converted to RGB, so that the pixel tensor shape matched the network’s expected input.

After this, the standard ImageNet normalization (subtracting the dataset mean and dividing by the standard deviation for each channel) was applied via the model’s preprocessing function, which centers pixel values and helps the network train more stable ^13^. To expand the effective dataset and reduce overfitting, on-the-fly data augmentation was incorporated: random horizontal and vertical flips, small rotations up to ±20°, random zoom and shear transforms. These augmentations simulate real-world variability—such as larvae in different orientations or under uneven illumination—so the model learns robust, invariant features, and their predictions are consistent irrespective of the positioning or lighting of the larvae.

#### Data Splitting and Label Encoding

Each class folder is split into a stratified training (85%), validation (15%), preserving class balance so that no single life stage would dominate the loss function. Each model was trained. Finally, our categorical labels were encoded with one-hot vectors for the eight stages, ensuring the softmax head could compute a cross-entropy loss directly. Altogether, this preprocessing regimen standardized our inputs, enriched diversity with realistic perturbations, and prepared our tensors for high-performance deep learning.

#### HyperParemeter Tuning and Training Strategy

To optimize model performance, several hyperparameters were evaluated, and it was found that a batch size of 64 struck the best balance between gradient estimate stability and GPU memory utilization. Moreover, a layer-freezing schedule was introduced for some models, keeping all pretrained weights frozen for the first 10 epochs, then unfreezing the top 40 layers for an additional 15 or so epochs (Table 1: Model Training) to gradually adapt high-level features while minimizing catastrophic forgetting ^14^.

**Table 1.** Model Training

|  | **InceptionV3** | **Resnet50** | **ConvNeXt** |
| --- | --- | --- | --- |
| **Image size (pixels)** | 299 x 299 | 224 x 224 | 224 x 224 |
| **Batch size** | 64 | 64 | 64 |
| **Optimizer** | Adam with learning rate of 0.0001 | Adam with learning rate of 0.0001 (Initial training) 0.00005 (Fine tuning phase) | Adam with learning rate of 0.0001 (Initial training) 0.00001 (Fine tuning phase) |
| **Loss Function** | Categorical Crossentropy | Categorical Crossentropy | Categorical Crossentropy |
| **Epochs** | 25 (early stop at epoch 18) | Initial epochs = 10<br>Fine tuning epochs = 15 | Initial epochs = 8<br>Fine tuning epochs = 18 |

**Table 2.**
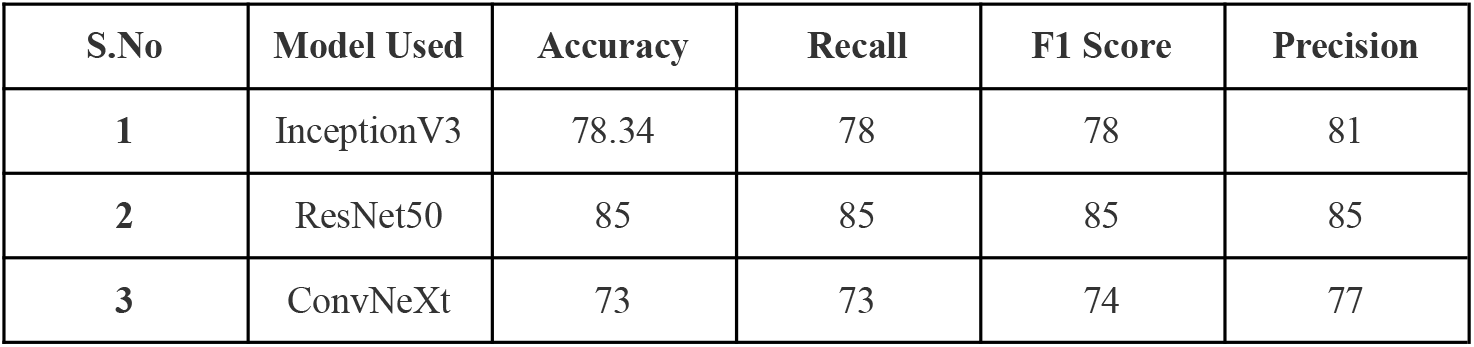
Model Performance

Furthermore, a lower learning rate of 1e-4 was employed to ensure smooth convergence and prevent the optimizer from overshooting optimal minima. During fine-tuning, the rate was decreased to 5e-5 to allow more delicate weight adjustments, helping the model refine high-level features. After training, the models were stored on Hugging Face for global deployment via the web app.

### Model Architectures

#### 1. ResNet-50

ResNet-50 employs a deep residual design where identity and convolutional blocks facilitate the training of very deep networks by preserving gradient flow ^10^. In the context of *Drosophila* stage detection, ResNet-50 was initialized with ImageNet-pretrained weights and modified for eight-class classification. All layers up to the 140th were frozen to retain general visual features, then a Global Average Pooling layer distilled spatial maps into feature vectors. Finally, a dense softmax head with eight outputs was appended to predict each developmental stage—from egg through black pupa—using a multi-class cross-entropy loss.

#### 2. InceptionV3

InceptionV3’s modular “Inception” blocks run convolutions of varied kernel sizes in parallel, capturing image details at multiple scales. For *Drosophila* stage detection, the network was loaded with ImageNet weights and adapted to accept 299×299 px inputs matching its native resolution ^10^. A new classification head (Global Average Pooling → Dense → Softmax) replaced the original, enabling the model to learn stage-specific morphological patterns ^11^.

#### 3. ConvNeXt

ConvNeXt reimagines classic convolutional networks with design elements inspired by vision transformers—such as inverted bottleneck blocks, large kernels, and simplified LayerNorm—yielding transformer-level accuracy at lower compute cost ^12,15^. In this project, ConvNeXtSmall was initialized with ImageNet weights and configured for 224×224 px inputs. The entire base was frozen during an initial training phase to adjust only the lightweight classification head (Dropout → Dense → Softmax). In a subsequent fine-tuning stage, all layers were unfrozen to specialize the model’s deep features for distinguishing the eight *Drosophila* life stages.

### Model Training

This study investigates the development and optimization of three state-of-the-art deep learning architectures for image classification tasks—ResNet50, InceptionV3, and ConvNeXt. Each model was systematically fine-tuned using tailored hyperparameter configurations, incorporating advanced data augmentation techniques and optimization strategies to maximize performance across diverse image inputs.

#### Staged Fine-Tuning with Reduced Learning Rate

We use a two-stage fine-tuning regime to adapt pretrained backbones without erasing their general visual features (Table 1: Model Training).

1. Stage 1 (Feature alignment, Higher learning rate). Freeze most (or all) of the backbone and train the classifier head at learning rate = 1e-4. This lets the model re-map high-level features to the new label space while keeping generic convolutional features intact.
2. Stage 2 (Backbone adaptation, Lower learning rate). Unfreeze the upper portion of the backbone (last ∼40 layers) in a single step and fine-tune with a reduced learning rate of 5×10^−5^ applied uniformly to both the unfrozen backbone and the classification head ^16^. This conservative schedule—with BatchNorm kept in inference mode (training=False), plus ReduceLROnPlateau and early stopping—helps stabilize statistics, limit feature drift, and mitigate catastrophic forgetting while maintaining a balanced precision–recall profile, especially for adjacent/ambiguous stages ^17^.

### Mobile Deployment

To allow other users to use the ML models for their work, a Streamlit web-app was developed that was hosted on Streamlit Cloud for global deployment. During the making, the Streamlit app was kept deliberately lightweight and adaptive so it would run smoothly on a phone’s browser without any extra wrappers or containers. The app relied on Streamlit’s built-in responsive layout—just a centered title, subheader, and full-width video component—so elements automatically resize to the screen’s width. To avoid re-downloading our InceptionV3 weights on each refresh, @st.cache_resource was used for the model loader, which ensures the heavy download happens just once per session.

For live camera input, streamlit-webrtc ^18^ was integrated with async_processing=True, allowing frame inference to run on a background thread; slower mobile CPUs simply drop unprocessed frames rather than freezing the UI. Moreover, some frames were skipped for their prediction to make the frame rate smoother. Together, these choices—caching, auto-scaling, asynchronous frame handling, and lightweight logic—deliver a responsive, mobile-friendly experience without additional native code or REST endpoints.

## RESULTS AND DISCUSSION

We evaluated all models using a stratified 15% hold-out split to preserve per-stage balance ^14^. Metrics were computed with Scikit-Learn ^19^. Accuracy (correct/total) was used as the global indicator ^20^, while recall (stage-specific sensitivity) captured how reliably each *Drosophila* stage was detected. Precision reflected how often predicted stages were correct, and the F1 score—the harmonic mean of precision and recall—summarized the precision–recall trade-off ^20^.

## Key findings

ResNet-50 tight metric band (85%) signals a flat error profile: false positives and false negatives are roughly in check, and precision is not being “bought” by sacrificing recall (or vice-versa). Under our stratified 15% hold-out, this balance indicates the model is not leaning on head classes to inflate accuracy and is instead generalizing across stages—important for datasets where adjacent stages are visually not distinct thus easily confusable.

InceptionV3 is conservative, precision (81) > recall (78): it’s choosier—fewer false positives but more misses. So you get fewer false alarms, but you miss a few real cases. Pick this when false alarms are costly—e.g., each positive needs a human check. The conservative bias keeps the review queue cleaner, thus the small recall drop is often a fair trade.

ConvNeXt has Lowest accuracy (73%) with precision (77) > recall (73) suggests under-detection of harder stages (likely look-alike instars/pupal transitions). Not recommended without further tuning (thresholding, class-balanced loss, augmentation for underrepresented stages).

Operational takeaway: Since all models show precision ≥ recall, the main lever to move overall performance is boosting sensitivity on difficult/rare stages—ResNet-50 already balances this best in the current setting.

Many stages, like the instar 1 and instar 2 or brown pupa and eye pupa, appear quite similar (Figure: 3), making them difficult to distinguish even for trained experts. Combined with real-world variations in lighting, orientation, and background, this makes the task inherently challenging. Achieving accuracy beyond this point would require models to capture extremely fine details while staying robust.

**Figure 3.**
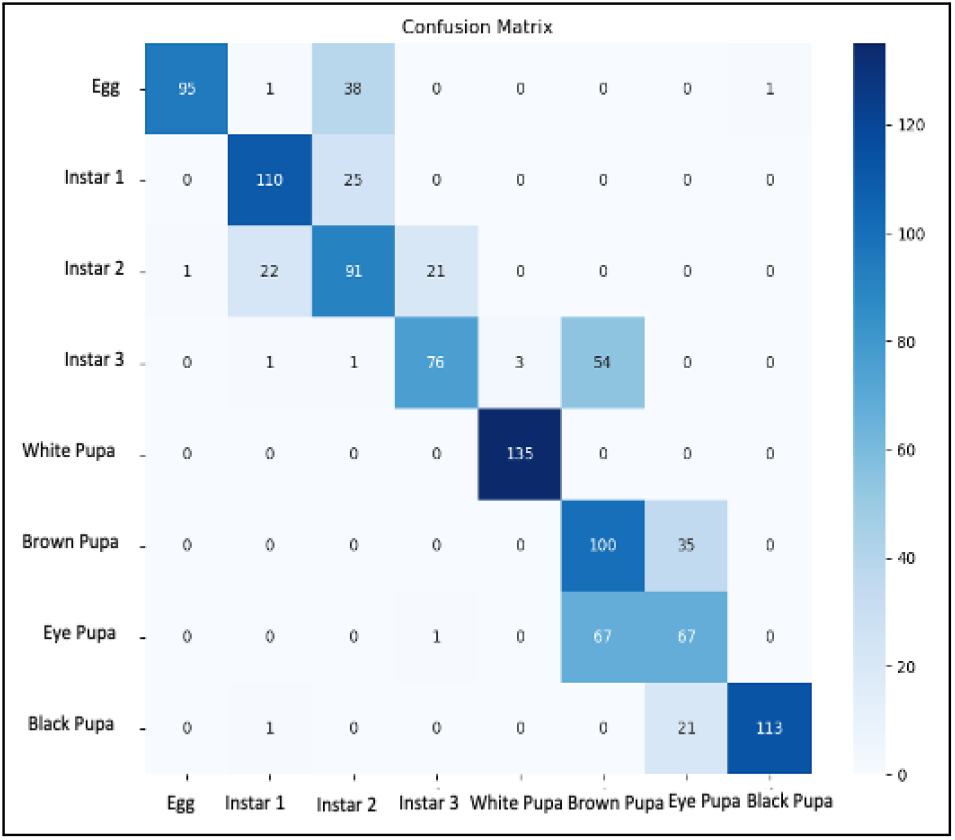
Confusion Matrix for *Drosophila* Life Stage Classification.

+The demonstrated performance has practical implications for biological and ecological research. First, achieving reliable stage classification enables high-throughput developmental studies without the bottleneck of manual annotation. This is particularly important in large-scale genetic or pharmacological screens, where thousands of individuals must be staged consistently. With an accuracy of 85%, it is better to use our web app than manually sorting through the *Drosophila*, as manually there may be bias which gives a greater error than our models. To remove biasness, multiple people must observe the same fly to come to a conclusion, that would be a waste of resource and time, this is precisely where these models come to use. Students, technicians, or researchers can rapidly stage samples at the bench or in teaching labs using only a camera-equipped device, lowering the barrier for adoption in diverse experimental workflows. By making the models, weights, and code publicly available, these results provide a scalable, community-driven tool that can be integrated into automated imaging pipelines or adapted to other insect models.

## CONCLUSION

This work presents a deep learning approach to automatically classify *Drosophila* life stages by fine-tuning multiple CNN models, including ResNet50, InceptionV3, and ConvNeXT. Standardized resizing, ImageNet-based normalization, and real-time data augmentation were applied to help the models learn robust features that generalize well under different lighting and poses. By using frame-skipping and asynchronous WebRTC processing ^18^, our system achieves real-time performance (∼8 FPS) with approximately 85% classification accuracy. This demonstrates that modern CNN architectures, when efficiently deployed via Streamlit, can reliably replace manual microscopy, offering a scalable approach to high-throughput phenotyping.

Future work may explore ensembling diverse backbones (say ViT or YOLOv8), integrating explainable AI modules to highlight discriminative morphological features, adding temporal modeling to smooth stage transitions, and packaging a standalone mobile app with on-device TensorFlow Lite inference for truly field-deployable phenotyping.

## SUPPLEMENTARY INFORMATION

Github Link:

1. Streamlit Webapp: https://github.com/RishiP2006/Drosophila_stages_identification_app
2. Model Training Script: https://github.com/RishiP2006/Drosophila_stages_models

Hugging Face link: https://huggingface.co/RishiPTrial/stage_modelv2

Web app link (For Mobile Use): https://drosophila-stages-identification-app.streamlit.app/

